# Tadpoles rely on mechanosensory stimuli for communication when visual capabilities are poor

**DOI:** 10.1101/2022.10.18.512729

**Authors:** Julie M. Butler, Jordan E. McKinney, Sarah C. Ludington, Moremi Mabogunje, Penelope Baker, Devraj Singh, Scott V. Edwards, Lauren A. O’Connell

## Abstract

The ways in which animals sense the world changes throughout development. For example, young of many species have limited visual capabilities, but still make social decisions, likely based on information gathered through other sensory modalities. Poison frog tadpoles display complex social behaviors that have been suggested to rely on vision despite a century of research indicating tadpoles have poorly-developed visual systems relative to adults. Alternatively, other sensory modalities, such as the lateral line system, are functional at hatching in frogs and may guide social decisions while other sensory systems mature. Here, we examined development of the mechanosensory lateral line and visual systems in tadpoles of the mimic poison frog (*Ranitomeya imitator)* that use vibrational begging displays to stimulate egg feeding from their mothers. *We found that tadpoles hatch with a fully developed lateral line system. While begging behavior increases with development, ablating the lateral line system inhibited begging in pre-metamorphic tadpoles, but not in metamorphic tadpoles.* We also found that the increase in begging and decrease in reliance on the lateral line co-occurs with increased retinal neural activity and gene expression associated with eye development. Using the neural tracer neurobiotin, we found that axonal innervations from the eye to the brain proliferate during metamorphosis, with little retinotectal connections in recently-hatched tadpoles. We then tested visual function in a phototaxis assay and found tadpoles prefer darker environments. The strength of this preference increased with developmental stage, but eyes were not required for this behavior, possibly indicating a role for the pineal gland. Together, these data suggest that tadpoles rely on different sensory modalities for social interactions across development and that the development of sensory systems in socially complex poison frog tadpoles is similar to that of other frog species.

## INTRODUCTION

Animals use a variety of sensory modalities to detect the environment around them. For species with parental care, the ability to detect and respond to a potential caregiver can be a matter of life-and-death. Caregiver recognition can be complicated by the fact that some sensory modalities continue to change throughout development, where young animals may need to make complex social decisions with limited or varying sensory capabilities. Investigating how animals recognize potential caregivers at an early developmental age and how this might change throughout development is critical for our broader understanding of how sensory systems develop and evolve for communication.

Many aquatic animals possess a mechanosensory lateral line system that allows them to detect near-by water movements (Dykgraaf 1933). The lateral line system is composed of neuromasts, or bundles of hair cells, similar to those in the mammalian inner ear. Nearby water movements create a shear force on the hair cells that opens mechanically gated ion channels, permitting the detection of hydrodynamic stimuli. In fishes, the lateral line system is important for rheotaxis (Montgomery, Baker, and Carton 1997), schooling (Partridge and Pitcher, 1980), prey detection (Schwalbe, Bassett, and Webb 2012), predator avoidance (Stewart et al. 2014; Canfield and Rose 1996), and social communication (Butler and Maruska 2016). While less is known about the role of the lateral line in tadpoles, it has been implicated in responding to water currents (Brown and Simmons 2016; Simmons, Costa, and Gerstein 2004) and predator avoidance (Jung, Serrano-Rojas, and Warkentin 2020). In many species of frogs, the lateral line system is nearly or fully developed at hatching (Lannoo 1987; Zelena 1964; Saccomanno et al. 2021; Jung, Serrano-Rojas, and Warkentin 2020; Roberts et al. 2009) but degrades during metamorphosis (Wright 1951; Zelena 1964; Brown and Simmons 2016). As such, mechanosensation via the lateral line system is a potential communication channel during early-life stages that could play a role in detecting and recognizing social stimuli.

Although the adults of many species use vision for detecting social stimuli, the young of many species have limited or absent visual capabilities. For example, cats and dogs are born with their eyes closed and are functionally blind, whereas mouse pups do not open their eyes until they are 11-12 days old (UCSF Lab Animal Resource Center). Although open or shut eyes are often used as a proxy for whether mammalian neonates can see or not, this assumption is less clear in young animals of other taxa, whose eyes remain visible throughout development. Since as early as the mid 1800s, frogs and toads have been fundamental in visual neuroscience research (for review, see Donner and Yovanovich 2020), as adults have a fully developed and complex retina similar to other vertebrates. Tadpoles of some species, however, hatch with externally visible, partially developed eyes, but their visual capabilities are not well-developed (Hoff et al. 1999). Grant et al. (1980) divided early development of the tadpole retina into four stages, with visual function being present at the end of the first stage shortly after hatching, but noted that a mature retina was not present until there is significant hindlimb development. In addition, retinal projections to brain regions important for visual processing, such as the optic tectum, are still forming throughout metamorphosis (Fujisawa 1987). Despite poor or absent vision, these young animals likely make complex social decisions by relying on information from other sensory modalities. However, it is unclear how retinal development coincides with development of other sensory modalities and the complex visual behaviors observed in tadpole behavioral ecology.

Poison frogs (Family Dendrobatidae) are an emblematic amphibian taxa known for their bright coloration, which advertises their chemical defenses to predators. These taxa display a wide range of parental care strategies, notably tadpole transport and egg provisioning (Summers and Earn 1999; Summers and Tumulty 2014; Weygoldt 2009). Tadpoles of these species also show an impressive diversity of complex behaviors, such as aggression and begging for egg meals from their parents by vigorously vibrating their bodies. Recently, it was hypothesized that poison frog tadpoles are dependent on visual cues for many of these social behaviors (Stynoski and Noble 2012; Fouilloux, Yovanovich, and Rojas 2022; Fouilloux et al. 2023). Indeed, research in the Strawberry poison frog (*Oophaga pumilio*), suggests that tadpoles use visual cues to recognize parents apart from heterospecifics (Stynoski and Noble 2012; Fouilloux, Yovanovich, and Rojas 2022) and sexually imprint on the color morph of their caregiver (Yang, Servedio, and Richards-Zawacki 2019). In dendrobatid frogs with extended parental care, parents dive into the small aquatic nursery during visits, which could generate water movements that stimulate the tadpole lateral line. Their movements on nearby fronds above the nursery might also create vibrations that are transmitted to the water and detectable by the lateral line. Despite the growing utility of poison frogs as a model system for neuroethology research, sensory system development in poison frog tadpoles has not been well studied.

Here we examined the development of the lateral line and visual systems in tadpoles of the mimic poison frog (*Ranitomeya imitator*). In this species, fathers transport hatched tadpoles to individual pools in bromeliad leaves. Every few days, the parents will visit the tadpole, and the tadpole will beg for food by rapidly vibrating its body (Yoshioka, Meeks, and Summers 2016). Tadpoles must also attempt to avoid predation from spiders and other tadpoles, as sympatric *Ranitomeya variabilis* tadpoles cannibalize *R. imitator* tadpoles (Brown, Morales, and Summers 2008). In *Oophaga pumilio* tadpoles, predation accounts for as much as 67% of tadpole mortality, emphasizing the importance of detecting and avoiding potential predators (Maple 2002). As such, *R. imitator* tadpoles may make complex decisions about the identity of pool visitors. Here, using anatomical, experimental, and transcriptomic approaches, we tested two hypotheses: (1) that tadpoles, independent of developmental stage, use mechanosensory stimuli detected via the lateral line system for decisions about begging, and (2) that vision facilitates begging behavior, but that young tadpoles have overall poor visual capabilities. Answers to these questions will help clarify the mechanisms by which *R. imitator* tadpoles process the diverse sensory inputs they receive during development and therefore better inform our understanding of behavioral mechanisms in tropical tadpoles.

## METHODS

### Experimental Animals

All *Ranitomeya imitator* tadpoles were captive bred in our poison frog colony from adult breeding pairs using standard animal procedures in our laboratory. Briefly, reproductive male and female *R. imitator* were housed in a glass terrarium (12×12×18 inch) containing several water pools, greenery, and a moss-substrate floor. Water pools were checked regularly for deposited tadpoles. Transported tadpoles were housed individually in circular containers (5 cm diameter) in a large aquarium (5 gallon) maintained at 26-28°C with constant recirculation. Tadpoles were fed brine shrimp flakes or tadpole pellets (Josh’s Frogs, Owosso, MI, USA) three times weekly. Tadpoles have access to sphagnum moss and tadpole tea leaves as extra sources of nutrients. We used adult *R. imitator* females from actively reproducing pairs as stimulus animals. All procedures were approved by Harvard University Animal Care and Use Committee (protocol #17-02-293) and Stanford Administrative Panel on Laboratory Animal Care (protocol #33097).

### Tadpole Development

All tadpoles used in behavior trials were measured for body mass, body length (mouth to tail peduncle), and total length. We developed a *R. imitator* staging guide based on Gosner staging (**Supplementary Doc**; (Gosner 1960) and a numerical stage (based on hindlimb development) was recorded for all tadpoles. When appropriate, we grouped tadpoles based on stage. Early-stage tadpoles had minimal pigmentation, no hindlimb development, and consisted of only stage 25 tadpoles. Middle-stage tadpoles were partially pigmented with a tan/gray coloring, had minimal hindlimb development (<1 mm), and consisted of tadpoles between stages 26-29. Late-stage tadpoles had full pigmentation, including some adult-typical pigmentation, significant hindlimb development, forelimbs had not emerged, and ranged from stage 30-40. Although this range of stages is large and encompasses many major metamorphic events, such a classification was necessary due to timing of experiments, feeding, and sample sizes.

### Begging Behavior Trials and Analysis

To examine begging behaviors, we exposed naive tadpoles of various developmental stages to reproductive females. All tadpoles were food restricted for 48-72 hours prior to begging assays. On the morning of the trial (8-10 AM), tadpoles were placed into a circular arena (5 cm diameter, 10 cm height) filled with 100 mL of prewarmed frog water or square acrylic arena (5×5×5 cm) filled with 45 ml of frog water and allowed to acclimate for 10 minutes. Arenas were placed on an LED lightpad and imaged from above using a GoPro camera. After an acclimation period, we recorded a 10 min baseline for each tadpole when no stimulus was present. We then added an *R. imitator* female and allowed them to interact for 20 minutes. Based on live observations, each tadpole was assigned as begging (at least two bouts of begging during 20 min trial) or non-begging (no begging during 20 min trial). To assess how behavior changes across development, 10 tadpoles were tested once a week for 6 consecutive weeks in begging trials. For all other experiments, early, middle, and late-stage tadpoles were only used once in behavior and immediately collected.

Videos were scored using BORIS (Friard and Gamba 2016). We quantified the number of bouts and time tadpoles spent begging, swimming or moving. Swimming behavior was quantified as the amount of time a tadpole spent swimming around the arena and had to involve multiple back and forth tail movements. In contrast, a “movement” was quantified when a tadpole performed a single tail flick to change position in the arena and was included in activity measures as 0.5 sec. Begging behavior involves the rapid vibration of the body/tail, often with the tail straight, and is performed with the tadpole at a >45° angle to the female (Summers and Earn 1999; Summers and Tumulty 2014; Weygoldt 2009).

### Lateral Line Characterization and ablation

To visualize the lateral line system in early and middle stage tadpoles, we immersed live tadpoles in 0.1% DASPEI solution prepared in frog water for 5 minutes in the dark. Tadpoles were anesthetized in buffered 0.01% MS-222 in frog water prior to imaging. Tadpoles were kept in the dark during staining and until the completion of imaging. Anesthetized tadpoles were placed in a small dish containing frog water and imaged on a Leica M165 FC stereo-dissecting scope equipped with Leica DFC 7000 T camera and a GFP filter. We imaged three very early (<2 days post transport), three early (1-2 weeks post-transport), and three middle stage tadpoles to create a consensus lateral line map for each developmental stage.

To test the role of the lateral line system in tadpole social behaviors, we chemically ablated the lateral line system using streptomycin sulfate (N=10 tadpoles per stage, per treatment). Tadpoles were immersed in 0.13% streptomycin prepared in 100 ml of frog water at room temperature for 1 hour the after afternoon before behavior trials. After treatment, tadpoles were allowed to recover in normal frog water overnight before begging behavior trials the next morning (as described above). Sham-treated tadpoles were handled as above, but the streptomycin was omitted from the treatment. To verify treatment efficacy, several tadpoles were stained with DASPEI as described above at the conclusion of behavior trials. Lateral line-ablated tadpoles had little-to-no remaining neuromasts, indicating that our ablation protocol is effective at functionally removing >90% of the lateral line and that these effects remain for at least 24 hours.

### Light Preference Trials

To assess tadpole visual capabilities and light environmental preferences across development, we developed a phototaxis assay for tadpoles (protocol is available on protocols.io dx.doi.org/10.17504/protocols.io.x54v9p294g3e/v1). We tested early, middle, and late-stage tadpoles in a light/dark preference assay (N=20-25 tadpoles per group). The behavioral arena was constructed from a 10 cm and 15 cm diameter. We burned a small hole for a bolt to pass through in the middle of the large petri dish and painted half of the bottom and sides of the larger petri dish black with multiple coats of black acrylic paint to ensure that no light passed through it. The other half was left unpainted. The dish was then sprayed with Kyrlon Fusion Clear Gloss to seal it from water and create a slight “frost” on the unpainted side that reduced reflections. A screw and bolt were used to attach the small petri dish to the lid of the large petri dish. The painted dish was placed between the two, with the screw going through a hole burned in the larger dish, so that the light/dark dish could be rotated. This setup allows us to change the light environment (flip the light/dark sides) without disrupting the tadpole in the small petri dish. The behavioral setup was placed on an LED light pad set to full power and a GoPro camera was used to record from above.

On the morning of the trials, tadpole containers were moved to a procedure room and allowed to acclimate to the room for ∼10 min. The small petri dish (tadpole arena) was filled with 40 ml of frog water. Tadpoles were transferred to the middle of the arena and behaviors were recorded for 3 minutes. We then flipped the light arena and recorded for an additional 3 minutes. A subset of tadpoles (N=9 per group) were used in the light preference tests on two consecutive days, with eye removal and neurobiotin labeling (see below for details) immediately following the first trial.

We scored each video for time spent in each light environment, as well the amount of activity (swim duration and number of movements) that occurred on each side of the arena. We also measured the latency to activity and the latency to enter the dark side of the arena. Since most tadpoles settled onto the dark side of the arena early in the trial, we also recorded if the tadpole “tracked” to the dark side of the arena after the sides were flipped and used this as an indication of a true preference being displayed.

### Neurobiotin Labeling

To examine when connections between the retina and brain are established in *R. imitator*, we placed neurobiotin into the cut optic nerve of early, middle, and late-stage tadpoles (N=9 tadpoles per group). This allowed us to visualize the optic nerve and its projections throughout the brain using fluorescent microscopy. Following the light preference trial, each tadpole was anesthetized in 0.01% MS-222 in frog water and the eyes were removed. Approximately 0.1 μl of 10% neurobiotin solution was placed in the eye cup onto the optic nerve. Tadpoles were placed into recovery containers in fresh frog water and allowed to recover overnight. The following morning, blinded tadpoles were run through the light preference trials before being collected. All blinded tadpoles recovered by the following morning, had normal swimming behavior, and responded to a water puff.

### Tissue Collection

We collected eyes from begging, non-begging, and control tadpoles to examine neural activation of the retina and molecular profiles associated with begging across development. For tadpoles used in begging trials, the female was quickly removed from the arena at the end of the trial, the light turned off, and the tadpole incubated in the arena for 30 min to allow for production of neural activation associated with the preceding begging trial. For tadpoles used for neurobiotin tracing, tadpoles were euthanized immediately after the second light preference test. To collect brains for immunohistochemistry, tadpoles were then euthanized with an overdose of benzocaine, the brain exposed, and the whole head fixed in 4% paraformaldehyde (PFA) prepared in 1x phosphate buffered saline (PBS) at 4°C overnight. The tadpole head was then rinsed in 1x PBS for 24 h, and cryoprotected in 30% sucrose prepared in 1x PBS at 4°C. Once dehydrated, tadpoles were embedded in OCT mounting media (Tissue-Tek® O.C.T. Compound, Electron Microscopy Sciences, Hatfield, PA, USA), rapidly frozen, and stored at -80°C until cryosectioning. We sectioned the whole tadpole head, including the brain and eyes, at 14 μm and collected sections into 4 sets of alternate series of SuperFrost Plus microscopy slides (VWR International, Randor, PA, USA). Slides were allowed to dry completely and stored at -80°C until processing. For eyes collected for molecular profiling (see details below), the tadpole was euthanized as above, but the eyes were quickly removed from the head and immediately frozen in phosphoTRAP dissection buffer. Both eyes from three tadpoles were pooled together for each sample to obtain enough RNA for library preparation.

### Immunohistochemistry

To compare neural activation in the retina of begging and non-begging tadpoles (N=7-9 tadpoles per group), we stained cryosectioned tadpoles with an antibody to detect phosphorylated ribosomes (phosphorylated ribosomal subunit 6; pS6). Similar to immediate early genes, pS6 labels recently-activated neurons (Knight et al. 2012). Here, we compared neural activation patterns associated with begging across development stages. A western blot for pS6 in *R. imitator* tadpoles shows a band at the appropriate size, but this band is absent in protein samples treated with protein phosphatase 1, indicating it is specific for only phosphorylated S6 ribosomes (**Supplemental Info**). Moreover, in preliminary studies on the time course of pS6 in poison frogs we found that waiting 30-45 minutes after a stimulus or behavioral interaction is ideal for capturing neural activation associated with the preceding stimulus. Slides were allowed to come to room temperature, washed with 1x PBS 3×10min, blocked in 1x PBS with 5% normal goat serum (NGS) and 0.3% triton-X, incubated in 1:5000 rabbit anti-pS6 (Invitrogen; cat# 44-923G) prepared in blocking solution overnight at 4°C. The following morning, slides were washed with 1x PBS for 3×10 min, incubated in 1:200 AlexaFluor 488 goat anti-rabbit secondary antibody prepared in 1x PBS with 5% NGS for 2 hours, rinsed 1x PBS for 3×10 min, incubated in deionized water for 10 min, and coverslipped with DAPI hardset mounting media. All slides were stored flat at 4°C until imaging.

To detect neurobiotin in tadpole brains, slides of cryosectioned brains were reacted with a fluorescently-labeled streptavidin. Slides were brought to room temperature, washed with 1x PBS 3×10 min, and treated with streptavidin (1:400 prepared in 1x PBS; Streptavidin, Alexa Fluor 568 conjugate; Invitrogen, S11226) for 1 hour in the dark. Slides were then washed in 1x PBS for 3×5min, rinsed in DI water, and coverslipped with DAPI hardset mounting media. Slides were allowed to dry at room temperature in the dark for 1 hour, sealed with clear nail polish, and stored at 4°C until imaging.

### Microscopy and Cell Counts

To visualize pS6-labeled neurons as a proxy for neural activation, stained eye sections were imaged at 20x resolution on a Leica DM6B microscope using a DFC9000 digital camera controlled by LASX software. All images were taken with approximately the same exposure and intensity settings. To quantify the number of labeled cells, images were loaded into FIJI image analysis software (Schindelin et al. 2012). Cell counts were done in a single eye for each tadpole. We quantified sections immediately adjacent to the optic disk (± 5 sections) for a total of 6-8 sections per tadpole. pS6-immunopositive cells were visible in both the ganglion cell layer and inner nuclear layer, so each layer was quantified separately. The inner nuclear layer likely included multiple cell types, including amacrine cell, bipolar cells, and horizontal cells, and was roughly separated into a middle and top sub-section. For each image, the region of interest was circled, the area quantified, and the number of pS6-labeled cells in each region was quantified.

To measure the amount of neurobiotin present in the tecum as a proxy for retinotectal projection density, we imaged tadpole brains as described above, with the same exposure and intensity settings used for all animals. Only one hemisphere of the entire tectum was imaged. We used the threshold function in FIJI to highlight pixels representing neurobiotin staining. Threshold was based on the background levels of each image. Once the pixels were highlighted, we used FIJI to measure the percent of the region of interest highlighted to calculate a “relative projection density”. This method controls for differences in tectal size/thickness. We also completed these measurements on control tadpoles that were blinded by removing their eyes (as described above) but did not have neurobiotin applied to the optic nerve. The relative projection density was below 1% for all controls. As tectal layers are not apparent until later developmental stages, we quantified the tectum in whole, instead of by layers. Tectal thickness was measured as the distance from the outside of the tectum to the inner edge along the optic ventricle. The two widest measurements were averaged together for each animal.

### Molecular Profiling of Active Neurons

We used phosphoTRAP to molecularly profile active neurons in the eye during begging; phosphoTRAP uses an antibody for the neural activation marker pS6 to purify transcripts being translated by phosphorylated ribosomes at the time of collection (Knight et al. 2012). RNAseq is then performed on the total (TOT) input RNA sample and the immunoprecipitated (IP) sample. RNA samples were processed and purified as previously described (Fischer et al. 2019; Knight et al. 2012). RNA samples were purified using a SMART-Seq v4 Ultra Low Input RNA Kit for Sequencing (Takara, Mountain View, CA, USA), followed by library preparation using the Nextera XT DNA Library Prep kit (Illumina, San Diego, CA, USA), both according to manufactures’ protocols. Pooled equimolar library samples were run on an Illumina Hi-Seq 2500. Sequencing results were aligned to a *R. imitator* transcriptome built from tadpole brain, eyes, and gut samples. Count data were generated using DESeq2 (Love, Huber, and Anders 2014) and analyzed in R using paired t-tests on each transcript between the TOT and IP samples, similar to that described in Tan et al. (2016; **Supplemental Info**). Fold changes were also calculated for each transcript as log2 (IP count / TOT count). Differentially expressed genes were defined as having a p-value under 0.05 and a fold change greater than 1.5 in either direction. We chose to use paired t-tests of transcripts within each group because this better reflects changes in expression associated with begging and reduces the variation present due to intrinsic variables (developmental stage, hunger, etc) that affect total count data. We also did not correct for multiple hypothesis testing in phosphoTRAP data due to the small sample size (N=3 per group) and large number of transcripts. Bonferroni corrections or similar procedures reduce statistical power and increase the chance of type II errors, especially in small sample sizes. Although these tests do reduce type I errors, their unacceptable effects on statistical power can hide potential biologically relevant results (Nakagawa 2004). A list of differentially expressed genes was generated for begging and non-begging tadpoles. Gene ontology terms were added using the GeneOntology Database (Ashburner et al. 2000; Gene Ontology Consortium et al. 2023).

### Statistical Analyses

All statistics were performed in R (v4.0.2). We used student’s t-tests and generalized linear mixed models (GLMM; Package: glmmTMB; Brooks, Kristensen, and Van Benthem 2017) to compare behaviors among groups, with animal ID as a repeated, random factor when appropriate. Similarly, pS6 data was analyzed using a GLMM with layer as a repeated factor and animal ID as a random factor. When appropriate, Tukey’s post-hoc tests were used to parse differences among groups. Correlations were done using Pearson correlations. All data were checked for statistical outliers prior to analyses and removed. The only outlier detected was in the begging group of pS6 staining in the retina, in which one individual had values three times higher than all other animals.

All graphs were produced in R using ggplot2 (Wickham 2016). Box plots are used throughout for data visualization. All data points are represented as closed circles, data mean as an ‘X’, and data median as a solid line. Boxes extend to the furthest data points within the 25/75th quartiles, and whiskers extend to the furthest data points in the 5/95th percentile. We used a volcano plot generated by ggplot2 for visualizing phosphoTRAP data. Data points were plotted as -log10(p-value) and log2(fold change) for each transcript. Lines as Y=1.3, X=-0.58, and X=0.58 represent significant cutoffs of P<0.05 and FC>1.5, respectively.

## RESULTS

### Older tadpoles are more likely to beg

To lay a foundation for understanding the sensory contributions to begging behavior, we conducted begging behavior assays with randomly selected tadpoles and reproductive females. We initially classified tadpoles as begging or non-begging independent of stage. In general, ∼70% of tadpoles begged during behavior trials (**Fig. 1A**), with an average begging duration of 85.652 sec (**Fig. 1B**). However, when we accounted for tadpole stage, there was uneven distribution of stage within each group (**Fig. 1C**). Begging tadpoles were more developed than non-begging tadpoles (P=0.003), suggesting age might impact the likelihood to beg. To examine this further, we tested tadpoles in begging trials weekly for 6 consecutive weeks. Tadpoles are more likely to beg after 3 weeks of development (**Fig. 1D**), which roughly corresponds to the transition from early (stage 25) to middle developmental stages. Although the number of begging bouts did not statistically differ, tadpoles spent more time begging to females during the trials occurring weeks 5-6 compared to those in weeks 1-3 (**Fig. 1E**; F_5,42_=2.644, P=0.036).

**Figure 1.**
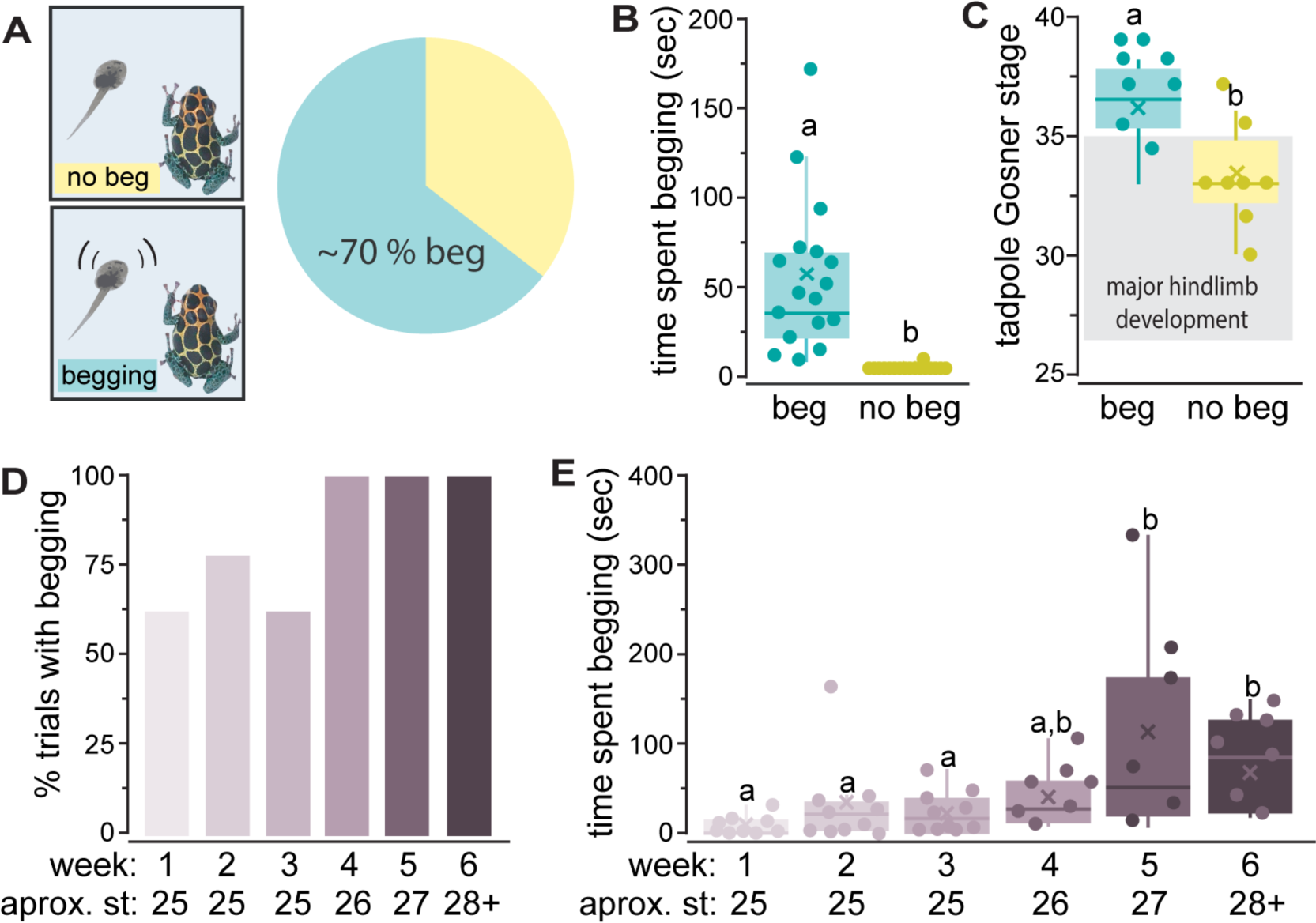
Tadpole begging increases during development. (**A**) When exposed to a reproductive female, ∼70% of tadpoles display begging behavior for an average time of 85.652 sec (**B**). (**C**) Begging tadpoles were of later developmental stages than non-begging tadpoles (P=0.003). (**D**) As tadpoles enter metamorphosis (st 26+; weeks 4-6), they are more likely to beg. (**E**) Older tadpoles spent more time begging (F_5,42_=2.644, P=0.036). Different lowercase and uppercase letters represent significant differences (P<0.05).

#### Tadpoles hatch with fully developed lateral line system

We first investigated if the lateral line system is fully developed at hatching in R. imitator tadpoles and if it mediates begging towards a potential caregiver. The lateral line system of R. imitator tadpoles is similar to that previously described in other anurans with several distinct lines of neuromasts and other clusters around the trunk and caudal peduncle (**Fig. 2A-D**) (Quinzio and Fabrezi 2014; Nieuwkoop 1975; Shelton 1971). An orbital line, with preorbital, supraorbital, and infraorbital portions, loops around the eye and extends to the medial side of the nares. On the dorsal side of the tadpole, the medial and dorsal lines extend down the body towards the caudal peduncle. A small cluster of neuromasts is located at the caudal peduncle. On the ventral surface of the tadpole, the oral line extends caudally from the sides of the mouth. The angular line extends from the side of the tadpole near the eye horizontally along the belly of the tadpole. The ventral line runs along the lateral portion of the belly and terminates at the caudal peduncle.

**Fig. 2.**
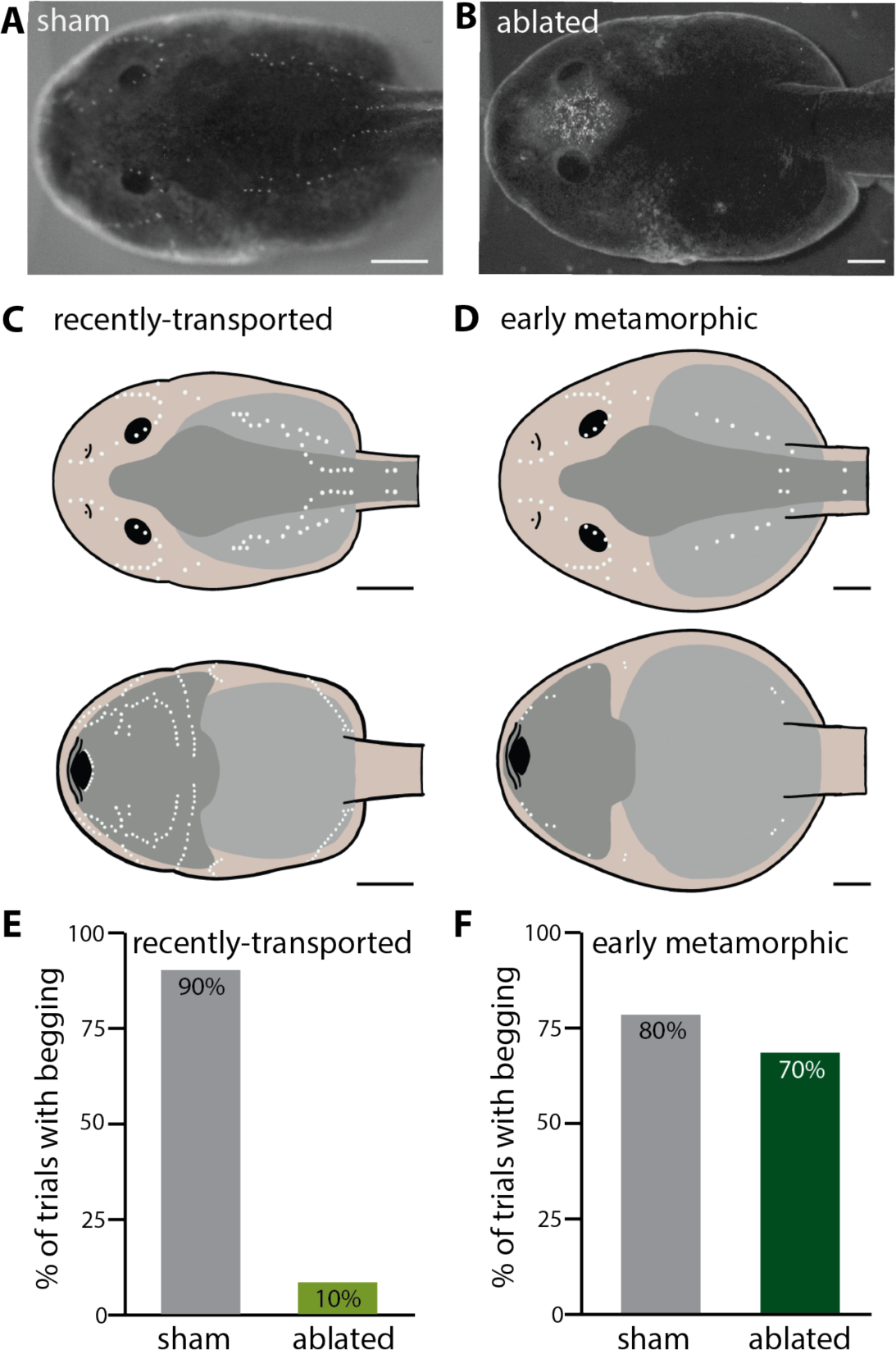
Lateral line system and use in behavior begging decreases with development. (**A-B**) Tadpoles were stained with DASPEI to visualize neuromasts in control (**A**) and streptomycin-treated (**B**) tadpoles. (**C-D**) Schematic of lateral line system in recently hatched (**C**) and early metamorphic (D) tadpoles. Neuromasts are depicted as white dots. Top and bottom schematics depict ventral and dorsal views, respectively. (**E-F**) Recently-transported tadpoles with an ablated lateral line system have reduced begging (**E**, *X^2^*=12.8, P<0.001), but ablating the lateral line did not impact begging in older tadpoles (**F**, *X^2^*=0.266, P=0.606). Scale bars represent 1mm.

DASPEI-labeled neuromasts were abundant on all tadpoles, with more neuromasts present on younger tadpoles than those in early metamorphosis (**Fig. 2C-D**). Tadpoles hatched within the last 48 hours had the most abundant neuromasts, with the most striking differences between very early and middle stage tadpoles. Very early and early stage tadpoles had prominent staining on both the ventral and dorsal sides of the tadpoles, but very few neuromasts were detected on the ventral side of middle-stage metamorphic tadpoles. The only prominent lateral line staining in middle-stage tadpoles was around the eyes in the preopercular lines.

#### Age-dependent effects of lateral line ablation on tadpole begging behavior

Tadpoles transported within the past 1-2 days (stage 25) displayed begging 90% of the time when presented with a potential caregiver, but ablating the lateral line system significantly reduced the likelihood of begging (10%; *X^2^*=12.8, P<0.001; **Fig. 2E**). In older tadpoles that have started metamorphosis, sham-treated and lateral line-ablated tadpoles begged with similar likelihood (sham: 80%; ablated: 70%; *X^2^*=0.266, P=0.606; **Fig. 2F**).

### Tadpole stage and begging impact retinal neural activation

Vision has been proposed to be important for begging behavior (Stynoski and Noble 2012; Fouilloux, Yovanovich, and Rojas 2022), so we quantified neural activation in the retina of begging and non-begging tadpoles. Begging tadpoles had higher activation in both the ganglion cell layer and inner nuclear layer compared to non-begging tadpoles (**Fig. 3A-B**; beg v no beg: F_1,15_=8.701, P=0.026; layer: F_2,15_=2.888, P=0.095). However, there was a significant positive correlation between tadpole stage and neural activation (R=0.557, P=0.025; **Fig. 3C**). To account for the developmental differences in begging and non-begging tadpoles, we also included a group of younger tadpoles from a separate study on begging and aggression. Within this group of stage 25 tadpoles, there was no statistical difference in neural activation in either the ganglion cell or inner nuclear layers among begging, aggressive, and control tadpoles (**Fig. 3D**, group:F_2,48_=0.422; P=0.658).

**Figure 3.**
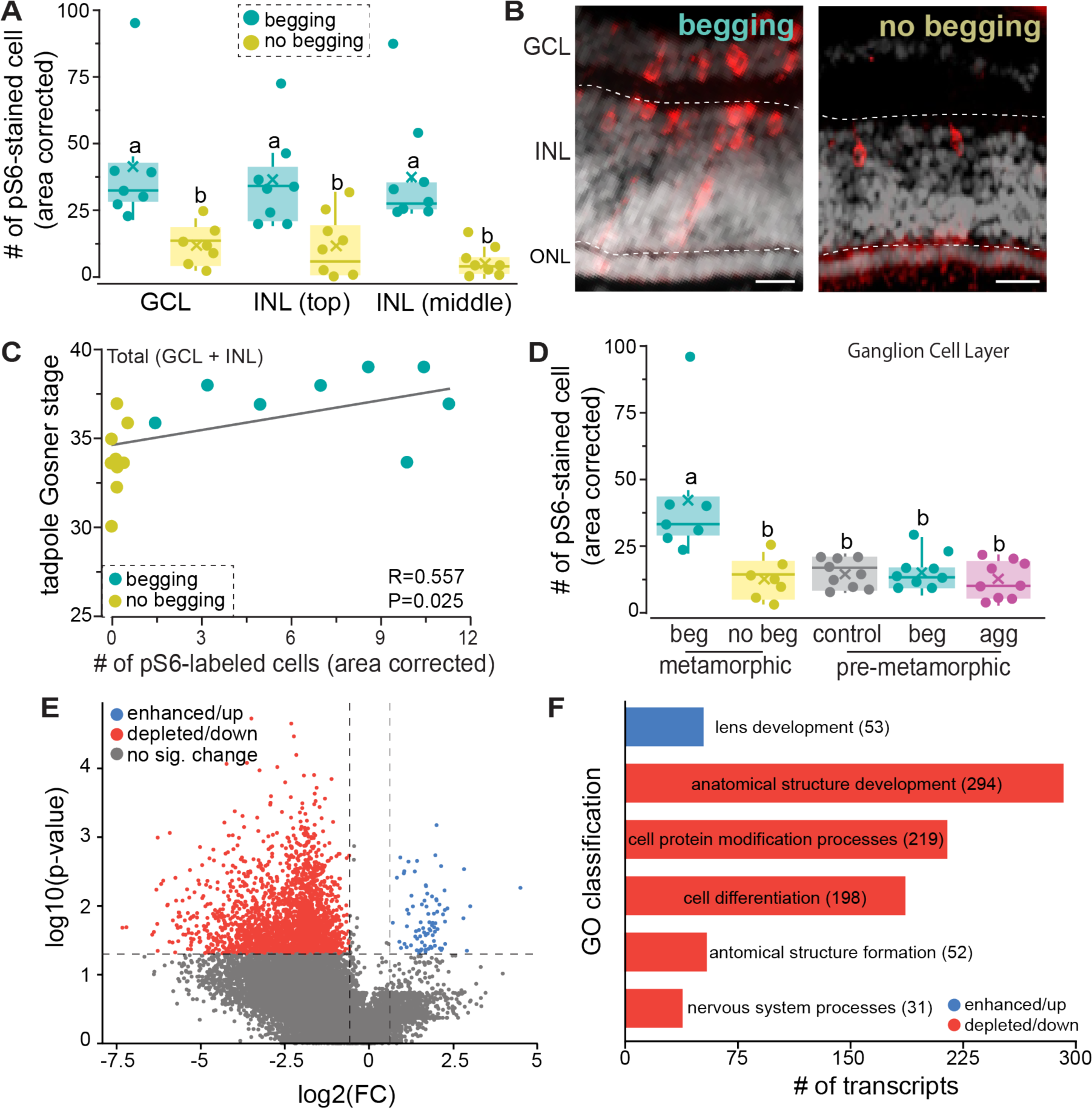
Retina neural activation varies with stage and begging. (**A**) Begging tadpoles had more pS6-immunopositive in the ganglion cell layer (GCL) and inner nuclear layer (INL) compared to non-begging tadpoles (F_1,15_=8.701, P=0.026). (**B**) Photomicrographs showing pS6-positive cells (red) in the GCL and INL, but not in the outer nuclear layer (ONL). Dotted lines approximately separate layers and gray is DAPI. Scale bar = 10 um. (**C**) The number of pS6-stained cells positively correlated to tadpole stage (R=0.557, P=0.025). (**D**) While metamorphic begging tadpoles had higher activation in the ganglion cell layer, there was no significant difference in pS6 staining in the retina of control, begging, and aggressive pre-metamorphic tadpoles (F_2,48_=0.422; P=0.658). (**E**) Volcano plot of phosophoTRAP data showing enhanced (blue) and depleted (red) transcripts in the eyes of begging tadpoles. (**F**) Gene ontology classification of enhanced and depleted transcripts reveals that over half of the enhanced transcripts are related to lens development, with cell development-related transcripts dominating depleted transcripts. The number of transcripts in each category follows the classification name. All differentially expressed genes and their GO terms can be found in the supplemental information. Different lowercase letters represent significant differences (P<0.05).

As older, begging tadpoles had higher neural activation in the retina, we next molecularly profiled these active neurons in eyes from begging and non-begging tadpoles using phosphoTRAP. There were 83 transcripts that were enhanced in neurons active during begging, but 2389 transcripts that were depleted in neurons active during begging (**Fig. 3E**). Of the transcripts that were up-regulated, 52 were *crystallin*-related, which is an important structural component of the lens (**Fig. 3F**). *Grifin*, a lens-specific protein, was also enhanced. Other potentially relevant up-regulated transcripts include several related to retinoic acid signaling and *sortilin*, an important regulator of neuron growth. Among the depleted or down-regulated transcripts associated with begging, there was a high number of transcripts associated with anatomical structural development (294 transcripts), cell protein modification processes (219 transcripts), cell differentiation (198 transcripts), anatomical structural formation (52 transcripts), and nervous system processes (31 transcripts). Another potentially important depleted transcript was *ephrin*, which is an important modulator of retinotectal connections and organization (McLaughlin et al. 2003). Despite more than 2400 transcripts enhanced or depleted in begging tadpoles, there were only 4 up-regulated and 18 down-regulated genes in the eyes of non-begging tadpoles. None of these genes overlapped between begging and non-begging tadpoles.

### Retinotectal connections increase during metamorphosis

To visualize connectivity between the retina and brain, we next applied neurobiotin, an anterograde neuronal tracer, to the optic nerve and quantified the amount of neurobiotin present in the tectum (**Fig. 4A, B**). This serves as a proxy for the extent of retinotectal connections and/or axonal branching in the optic tectum. Late stage tadpoles had more neurobiotin present in their tectum than younger tadpoles (F_2,21_=22.326; P<0.001), but middle stage tadpoles had more than early stage tadpoles (**Fig. 4C**). Relative projection density was positively correlated with developmental stage in late stage tadpoles (R=0.905; P<0.001; **Fig 4D**). Little to no fluorescence was observed in un-labeled tadpoles (average density <1%). Neurobiotin density was controlled for by tectum size, since older tadpoles had wider tectums than younger tadpoles (F_2,21_=88.476; P<0.001), and middle stage tadpoles as an intermediated between early and late stage tadpoles (**Fig. 4E**). Although not quantified, the lens of younger tadpoles appeared to have more DNA present (visualized via DAPI staining) than in older tadpoles, suggesting that it is still developing.

**Figure 4.**
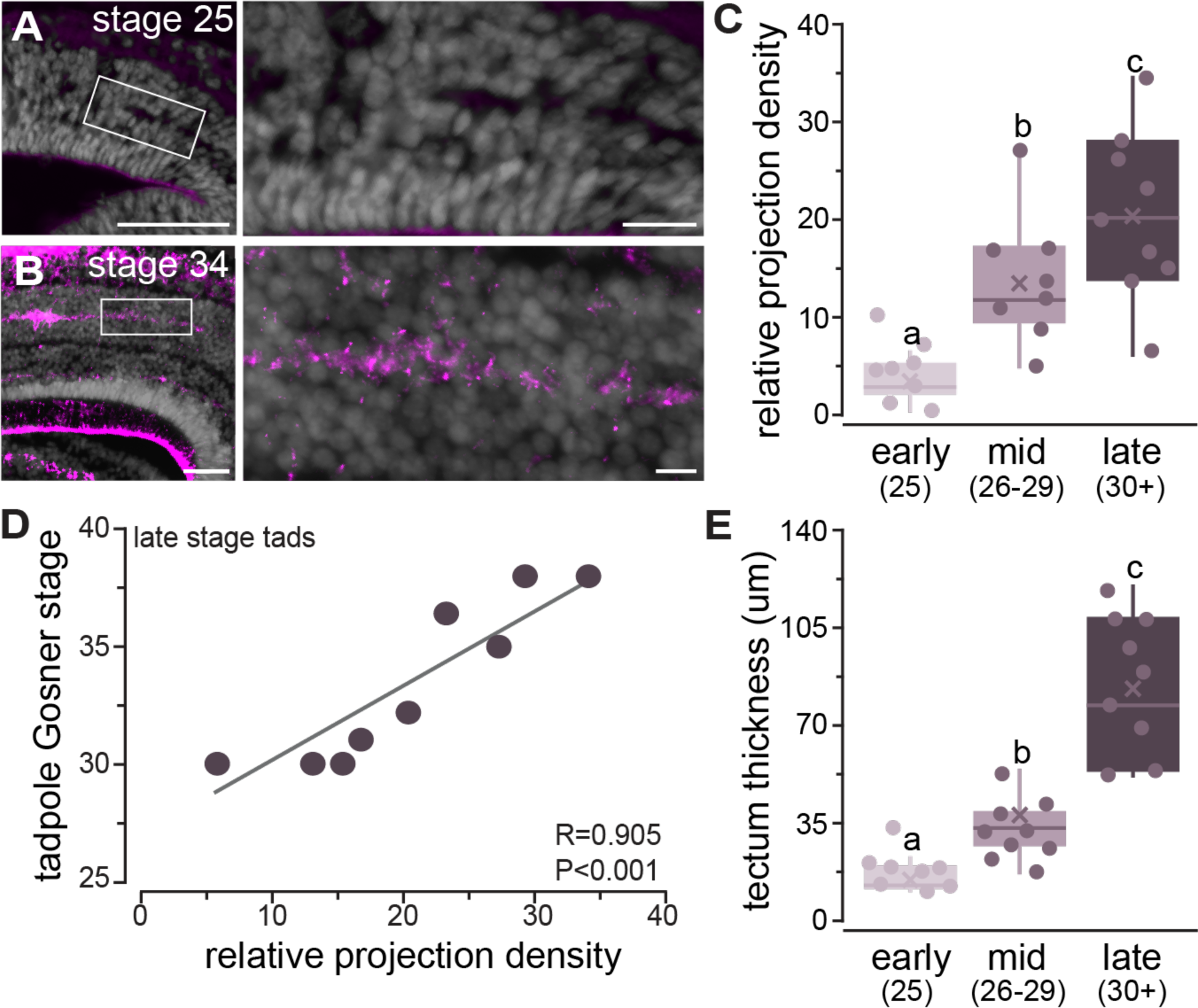
Retinotectal projections increase during metamorphosis. (**A-B**) Photomicrographs of fluorescently-detected neurobiotin in the tectum after it was applied to the optic nerve of an early (A) and late (B) stage tadpole. (**C**) More neurobiotin was detected in the tectum as tadpole stage increased (F_2,21_=22.326; P<0.001), indicating increased retinotectal projections and/or branching. (**D**) In late-stage tadpoles, stage positively correlates with the amount of retinotectal connections (R=0.905; P<0.001). (**E**) Thickness of the tectum also increased with tadpole stage (F_2,21_=88.476; P<0.001). Scale bars represent 50 um and 12.5 µm left and right images in A-B, respectively. Different lowercase letters represent significant differences (P<0.05). Numbers in parentheses in the x-axis of C represent the developmental stages included in each group, which are the same in E.

### Tadpoles prefer dark environments

Since we found differences in retinotectal projection densities with development, we next tested visual sensitivity in a light/dark preference assay. Early, middle, and late-stage tadpoles were tested in a light/dark preference arena (**Fig. 5A**), where the environment was flipped halfway through the trial to see if tadpoles tracked to one side over the other (**Fig. 5B**). All tadpoles, independent of stage, showed a preference for the dark side of the arena, as evident by the time spent on the dark side and the ability to track to the dark side when the arena was flipped (**Fig. 5C,D**). The strength of the preference increased with age, with late-stage tadpoles spending more time on the dark side than early and middle-stage tadpoles (F_2,68_=6.263; P=0.003). Furthermore, all late-stage tadpoles tracked to the dark side of the arena after the sides were flipped, but only 65% and 75% of early and middle stage tadpoles, respectively, displayed side tracking behavior. Late-stage tadpoles tracked to the dark side sooner than early and middle stage tadpoles (F_2,63_=3.707; P=0.030). When first entering the arena, late-stage tadpoles also entered the dark side sooner (F_2,68_=4.812; P=0.011) than younger tadpoles (**Fig. 5E**) and all late-stage tadpoles explored the dark side while only 81% of younger tadpoles initially explored the dark side of the arena. This cannot be explained by activity, as there is no significant difference in the total activity time (F_2,68_=1.436; P=0.245) among the different stages. In addition, the amount of activity on the dark side of the arena did not statistically change with developmental stage (F_2,68_=2.182; P=0.120). As such, the higher preference for the dark side displayed by older tadpoles is likely due to either increased light detection capabilities and/or increased motivation.

**Figure 5.**
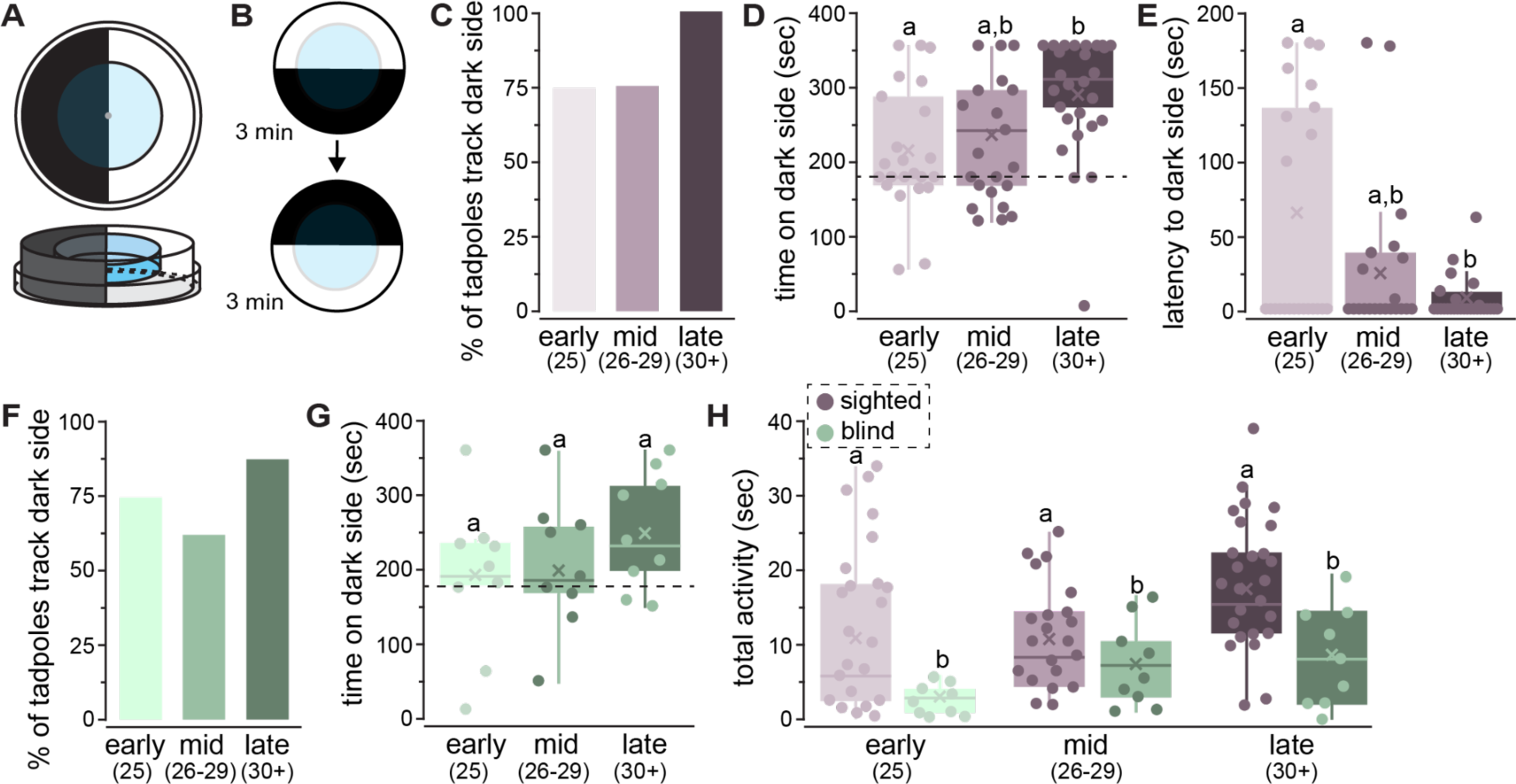
Older tadpoles show a stronger preference for dark environments, independent of sight. (**A-B**) A behavioral arena was constructed so that the light environment could be flipped halfway through the trial without touching the tadpole dish. (**C**) All late-stage tadpoles tracked to the dark side after the arena flip, but only ∼75% of younger tadpoles displayed this behavior. (**D-E**) Late-stage tadpoles spent more time in the dark environment (F_2,68_=6.263; P=0.003) and entered it earlier compared to early-stage tadpoles (F_2,68_=4.812; P=0.011). (**F-G**) Blind tadpoles also display a preference for the dark environment (F_2,23_=0.793; P=0.466). (**H**) Blind tadpoles were less active than sighted tadpoles (F_1,53_=8.196, P=0.021), but activity did not differ with stage (F_2,53_=3.418, P=0.058). Dotted lines in D and G are placed at 180 sec, or 50% of the total trial time. Different lowercase letters represent significant differences (P<0.05). Numbers in parentheses represent the approximate Gosner stage included in each group.

### Blinded tadpoles still show a dark preference

Tadpoles that were blinded for neurobiotin labeling still showed a preference for the dark environment (**Fig. 5F, G**). In total, 75% of blinded tadpoles displayed side tracking behavior after the arena was flipped, with late-stage tadpoles the most likely (88%) to track to the dark side. Even though blinded tadpoles were able to track to the dark light environment, their activity was dramatically reduced compared to sighted trials (**Fig. 5H**; blind v sight: F_1,53_=8.196, P=0.021; stage F_2,53_=3.418, P=0.058), but this decrease in activity did not vary significantly by stage (sight*stage interaction: F_2,53_=0.101, P=0.904). There were no significant differences among the three stages of blinded tadpoles for time spent in the dark environment (F_2,23_=0.793; P=0.466), latency to enter the dark environment (F_2,23_=1.477; P=0.251), or activity (F_2,23_=3.131; P=0.066).

## DISCUSSION

Animals rely on sensory information to carry out vital life processes, even at a young age. Visual capabilities are well documented to vary throughout life, where neonates of several taxa have absent or reduced vision. However, even young animals have to make complex decisions about their environment and may rely on other sensory modalities that are more developed. Here, we examined development of the lateral line and visual systems in *Ranitomeya imitato*r, a species of dendrobatid poison frog with complex social behaviors observed between parents and offspring (Summers and Tumulty 2014; Brown, Morales, and Summers 2008). We first hypothesized that tadpoles use mechanosensory stimuli to facilitate decisions about begging. We also hypothesized that vision facilitates begging behavior in a stage-dependent manner, with younger tadpoles having overall poor visual capabilities. We found that both visual and lateral line morphology changed throughout development, and that tadpoles rely on different sensory modalities for caregiver recognition throughout development, consistent with the ontogeny of the visual and mechanosensory systems.

### Begging behavior changes across development

Mimic poison frog tadpoles make crucial decisions with each visitor to their nursery, because they must assess each visitor to decide whether or not it is appropriate to beg for food. This process is required for feeding, but is also energetically costly (Yoshioka, Meeks, and Summers 2016; Stynoski, Stynoski, and Noble 2018), and begging to a non-caregiver may increase predation risk (Stynoski and Noble 2012). Strawberry poison frog (*Oophaga pumilio*) tadpoles need multimodal cues to elicit begging (Stynoski and Noble 2012), and Stynoski and Noble (2012) emphasized the importance of visual cues. In addition, *O. pumilio* tadpoles are thought to imprint on their parents’ coloration (Yang, Servedio, and Richards-Zawacki 2019), further suggesting visual cues are important for tadpole social recognition. In laboratory conditions, *R. imitator* tadpoles reliably beg to any reproductive female (∼75% in these experiments), but some tadpoles do not. This variation in begging was at least partially related to tadpole stage, since begging tadpoles were of a later developmental stage than those that did not beg. In addition, metamorphic tadpoles were more likely to beg than pre-metamorphic tadpoles. When metamorphic tadpoles did beg, they begged for a longer period of time than pre-metamorphic tadpoles, possibly increasing their chances of being fed (Yoshioka, Meeks, and Summers 2016). This result aligns with previous research showing that *R. imitator* tadpoles increased their begging with age (Brooks, Talbott-Swain, and Dugas 2023). It is important to note that the studies in *O. pumilio* were conducted using metamorphic tadpoles (Stynoski and Noble 2012: Gosner stages 30-40; Stynoski, Stynoski, and Noble 2018: stage 28+), when vision is likely more developed. In one previous study on *R. imitator* begging, the authors noted that they only used tadpoles >14 days post-hatch, because younger tadpoles did not display begging (Yoshioka, Meeks, and Summers 2016). A similar pattern of increased begging with age can be seen in many birds; for example, Kilner (2001) noted that “begging displays become increasingly flamboyant as chicks near independence.” House wrens (*Troglodytes aedon)* increase their begging with age, but the rate of increase is dependent on offspring fitness and brood size (Bowers et al. 2019). It is possible that the increase in begging observed in metamorphic tadpoles reflects higher nutritional needs associated with early metamorphosis (Pandian and Marian 1985).

#### Lateral line system

Consistent with what that described in other anurans, *R. imitator* tadpoles are transported with developed lateral line systems. *Xenopus* tadpoles just prior to and after hatching are able to detect water currents, and electron microscopy shows developed stereocilia and kinocilia (Roberts et al. 2009). In some amphibian species, like *Xenopus laevis*, neuromast morphology is modified during metamorphosis and retained through adulthood (Shelton 1970). While the visual system continues to develop throughout metamorphosis in *R. imitator* tadpoles, the number of neuromasts, the functional unit of the lateral line system, decreases over time, consistent with lateral line system degeneration during metamorphosis described in other frog species (Wright 1951; Zelena 1964; Brown and Simmons 2016). This is consistent with a decrease in reliance on the mechanosensation for begging behavior. Although begging generally increases throughout development, we have also observed that recently-transported tadpoles will reliably beg to a caregiver. After transport and deposition into their nursery, tadpoles are visited and fed by mom within the first couple of days. Ablating the lateral line system during this period almost completely inhibited begging (reduced from 90% to 10% of tadpoles), whereas ablating the lateral line system in older tadpoles had no impact on begging. Overall, this suggests tadpoles might rely on different sensory modalities throughout development for decisions about begging.

While the lateral line is known to be involved in fish social interactions, this is the first study, to our knowledge, to implicate it as important for tadpole affiliative social behaviors. In fishes, mechanosensation is used for detecting nearby individuals, assessing opponents, evaluating mates, and coordinating courtship rituals (Butler and Maruska 2016; Webb et al. 2021; TerMarsch and Ward 2020; Yoshizawa et al. 2014). In *Xenopus* tadpoles, the lateral line is involved in directional current detection as well as maintaining position or posture within a water column (Simmons, Costa, and Gerstein 2004). In pre-freeding stages of *Xenopus* tadpoles, the lateral line was shown to be important for responding to water jet and suction stimuli (Saccomanno et al. 2021). Red-eyed treefrog, *Agalychnis callidryas*, embryos hatch prematurely to escape egg-predators, which was shown to be dependent on the lateral line system in early developmental stages prior to vestibular system development (Jung, Serrano-Rojas, and Warkentin 2020). As such, the lateral line system in tadpoles has been shown to be important for detecting various types of water movements as well as mechanical stimuli associated with predator avoidance. For dendrobatid tadpoles, their lateral line is likely important for detecting water movements associated with their caregiver on a nearby frond or in their nursery. Although not tested, it is possible that the type or strength of vibration and water movements could provide information about body size and, thus, identity. As the lateral line is functional at hatching, it likely plays a crucial role in detecting potential caregivers prior to visual system maturation.

### Changes in eye structure and function across development

Unlike hatching with a fully developed lateral line system, tadpoles hatch with under-developed eyes and are thought to have poor vision (Hoff et al. 1999). Grant et al. (1981) divided early development of the tadpole retina into four stages, with visual function being present at the end of the first stage (pre-metamorphic) but noted that a mature retina was not present until the end of the metamorphosis. In *Xenopus* tadpoles, the eyes undergo the most pronounced morphological and physiological changes during early metamorphic stages. This developmental work can be juxtaposed with the behavioral ecology literature, where a recent study suggested that vision is important for poison frog tadpoles during social interactions (Fouilloux, Yovanovich, and Rojas 2022; Fouilloux et al. 2023), who often develop in water environments with low visibility. If vision is important for tadpole begging behaviors, we predicted that begging tadpoles would have higher neural activation in the retina compared to non-begging tadpoles. Although metamorphic begging tadpoles did have higher neural activation in both the ganglion cell layer and inner nuclear layer, pre-metamorphic begging, control, and aggressive tadpoles had similar levels of neural activation in the retina. The ganglion cell layer, which consists of neurons whose axons comprise the optic nerve, is mostly developed by tadpole hatching, whereas cells are continually added to the inner nuclear layer during tadpole development (Hollyfield 1968). Neuromodulatory amacrine cells in the inner nuclear layer do not develop until early metamorphosis, which coincides with the onset of expression of neuromodulators important for retina processing, such as tyrosine hydroxylase (synthesizes dopamine; Huang and Moody 1995; Sarthy et al. 1981; Reh and Tully 1986) and neuropeptide Y (Hiscock and Straznicky 1990; Huang and Moody 1995). In our study, neural activation in our older metamorphic tadpoles significantly correlated with developmental stage. One possibility is that the cellular mechanisms and/or circuitry leading to the higher activation in begging metamorphic tadpoles is simply not present in pre-metamorphic tadpoles. Future studies should examine the types of cells activated in the retina of begging metamorphic tadpoles compared to younger begging tadpoles.

We used phosphoTRAP to examine transcripts being actively translated in the eyes of begging and non-begging tadpoles. We initially expected that eyes of begging tadpoles would have enhanced expression of some behaviorally-relevant neuromodulatory transcripts. Although there were over 2400 enhanced and depleted transcripts in begging tadpoles, many of these were related to development. For example, begging tadpoles had a high number of enhanced transcripts associated with lens development. The increased expression of *crystallin-*related genes in the older group of tadpoles that displayed begging could reflect ongoing lens maturation. In *Xenopus* tadpoles, the size and density of the lens does not change during metamorphosis (Polansky and Bennett 1973), but the expression of *crystallins* does change across developmental stages (Polansky and Bennett 1973; Mizuno et al. 1999). Lens development in fish is largely complete before hatching, and it only grows in size as the animal develops and grows (Greiling and Clark 2009). However, ongoing lens development in *R. imitator* tadpoles is consistent with what has been observed in other tadpole species (e.g., Polansky and Bennett 1973). We also noted that older, begging tadpoles had a high number of depleted transcripts associated with development. This could reflect the developmental stage difference between begging and non-begging tadpoles, and suggests that tadpole stage, not the display of begging behavior, was driving our phosphoTRAP results. Overall, our data suggest that *R. imitator* tadpole eyes undergo morphological and physiological changes through late metamorphic stages, similar to that described in other frogs.

### Retinotectal projection development

We used the neural tracer neurobiotin to examine connections between the eye and the optic tectum. The optic tectum is the primary target of retinal ganglion cells in the eye, with immature connections between the eye and tectum forming within hours of fertilization in *Xenopus* tadpoles (Liu, Hamodi, and Pratt 2016). Some neurobiotin was detected in early-stage *R. imitator* tadpoles, indicating that connections between the eye and brain are present. Despite loading similar amounts of tracer into the optic nerve of early, middle, and late-stage tadpoles, the amount of tracer detected in the tectum increased with developmental stage in *R. imitator* tadpoles. This result suggests that retinotectal projections increase during metamorphosis, which is further supported by the correlation between the tracer abundance and developmental stage. One explanation for the increased neurobiotin observed in older tadpoles could be due to increased axonal branching and the establishment of circuitry within the tectum (Fujisawa 1987; Holt and Harris 1983; Fawcett 1981). In both *Xenopus* tadpoles and poison frog tadpoles, retinotectal projections rapidly expand during metamorphosis with only minimal retinotectal connections and axonal branching present at hatching. Overall, this suggests an increase in signals being sent from the retina to the brain associated with metamorphosis.

### Tadpole phototaxis behavior

We sought to test visual capabilities using a light/dark behavioral assay. Light preference assays are a common behavioral tool to assess exploratory, anxiety-like behaviors, and visual capabilities (Maximino et al. 2010; Takao and Miyakawa 2006). For example, rodents display pro-exploratory behaviors in novel environments, but also have an aversion to bright open areas (Bourin and Hascoët 2003; Shimada et al. 1995). When placed in the arena, most tadpoles initially chose one side and displayed some exploratory behaviors on that side. Midway through the trials, we flipped the light environment to test if tadpoles would track to the same light environment, indicating a true preference. In general, all tadpoles displayed a preference for the dark environment. The strength of this preference was strongest in late-stage tadpoles that all tracked the dark environment and spent more time on and entered the dark side sooner. In tadpoles, light preference and phototactic behaviors seem to vary with species. *Xenopus* tadpoles show a preference for the white side of a white/black preference assay, but when the optic nerves are severed, this preference goes away (Viczian and Zuber 2014; Moriya et al. 1996). This preference is dependent on both developmental stage and light environment during development. For example, Moriya et al. (1996) found that young *Xenopus* tadpoles show a preference for the white environment, but metamorphic tadpoles lose this preference, and froglets all display a preference for black (Moriya et al. 1996). The preference for light in *Xenopus* has been tied to retinal photosensors (type II opsins), with their expression increasing during development (Bertolesi et al. 2021). Generally, tadpoles display a preference for brighter light environments because they are often more nutrient rich (Jaeger and Hailman 1976), but some species may choose a light environment that allows them to be inconspicuous and thus reduces predation risk (Eterovick et al. 2020; Bertolesi et al. 2021). Since *R. imitator* tadpoles are raised in small pools in bromeliad leaves, predator avoidance, especially from above, may be the dominant motivating factor for the observed dark side preference. However, this idea needs to be tested in future experiments to investigate if the presence of a predator changes light environment preferences.

We found that eyes were not needed for a phototactic response in *R. imitator* tadpoles. In *Xenopus* tadpoles, a phototactic response towards the light side has been attributed to the photosensitive cells in the pineal structure (Foster and Roberts 1982; Mrosovsky and Tress 1966; Adler 1976). However, severing the optic nerve in some studies removed the preference for the white background environment (Viczian and Zuber 2014). It is important to note that these studies used a white/black assay, not a light/dark assay. Although a black arena will have lower light levels than the neighboring white side, the difference in illumination between the two sides is much greater in a light/dark assay that uses backlighting. This subtle difference could explain why eyes are important in some behavioral conditions, but not others, although, to our knowledge, this difference has not been experimentally tested. Light levels detected via the pineal structure are also involved in swimming behaviors in *Xenopus* (Jamieson and Roberts 1999, 2000; Li, Wagner, and Porter 2014). In our experiments, even blinded tadpoles displayed a behavioral preference for the dark side of the arena, indicating eyes are not needed for their phototactic response. However, blinding tadpoles removed the stronger dark side preference exhibited in late-stage tadpoles, suggesting that eyes likely play some role in light detection in late-stage tadpoles, consistent with our results that retinotectal projections increase during this time. The role of the pineal in detecting light is especially interesting given the natural behaviors of these poison frog tadpoles, where individual young are reared in small pools in bromeliad leaves. The parents visit the tadpole in its nursery, often approaching from above. This likely creates some sort of shadow that the tadpoles can detect, potentially via the pineal structure, as has been observed in blind cavefish (Yoshizawa and Jeffery 2008). Unfortunately, pinealectomies are difficult in *R. imitator* tadpoles, as there are no visually identifying structures of the pineal gland. Thus, the role of pineal structure on poison frog tadpole phototactic behavior remains to be tested. This result also emphasizes that, although vision is likely important for tadpole behavior, they rely on multisensory information from their environment (Stynoski and Noble 2012; Rot-Nikcevic, Taylor, and Wassersug 2006; Saidapur et al. 2009). Non-visual information may be especially important in young tadpoles with poorly developed visual systems relative to adults or in animals that live in light-limited environments.

## Conclusions

We show that poison frog tadpole sensory systems change throughout development, with a decreasing reliance on mechanosensory stimuli during social interactions coinciding with an increase in visual capabilities. Full visual capabilities likely do not emerge until metamorphosis, as suggested by the presence of development genes in phosphoTRAP data, changes in retina morphology, and retinotectal connections. Despite an underdeveloped visual system, even young tadpoles display begging behaviors towards potential caregivers, potentially based on vibrational stimuli detected via the lateral line system. Although not tested here, tadpoles likely rely on other sensory systems, like olfaction, for caregiver recognition and future work should focus on how tadpoles use multimodal cues for behavioral decisions. Together, this work emphasizes the importance of examining sensory system development in young and the role of multisensory interactions in social decision-making.

## ACKNOWLEDGEMENTS

We thank members of the O’Connell and Edwards labs, especially Madison Lacey, David Ramirez, Mesi Fisher, and Billie Goolsby, for their assistance with animal care and discussions about this research. We thank the Harvard University FAS Bauer Core for facilitating the transcriptomic work. We also thank Dr. Eva Fischer for discussions about analyzing phosphoTRAP data.

## DATA STATEMENT

All data associated with phosphoTRAP has been uploaded as supplemental information or to data repositories. The transcriptome will be uploaded to Dryad upon acceptance. Other data will be uploaded upon acceptance. The phototaxis assay is available on protocols.io at dx.doi.org/10.17504/protocols.io.x54v9p294g3e/v1.

