## Supplemental Information for "Tadpoles rely on mechanosensory stimuli for communication when visual capabilities are poor"

**Supplemental Files and Figures**

File 1: *Ranitomeya imitator* developmental staging guide for tadpoles from hatching through metamorphosis. (PDF file).

File 2: Video of light preference trial. A more detailed protocol for constructing arenas is found at dx.doi.org/10.17504/protocols.io.x54v9p294g3e/v1.

File 3: R script used for phosphoTRAP analyses. The *R. imitator* transcriptome will be uploaded to Dryad upon acceptance.

File 4: Count table generated during phosphoTRAP and used for analyses.

File 5: Excel file of all differentially expressed genes from phosphoTRAP analyses. Begging and non-begging, and up and down DEGs, are located on separate sheets.

File 6: Excel file of GO terms for down-regulated differentially expressed genes of begging tadpoles (corresponds to Fig. 3F).

File 7: *Ranitomeya imitator* tadpole ethogram (PDF file).

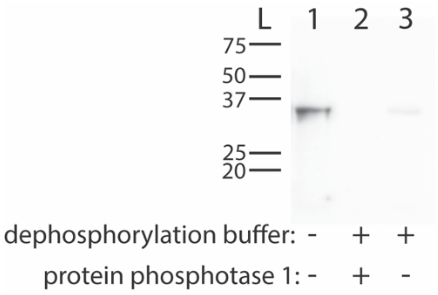


**Supp. Fig. 1. Western blot validating pS6 antibody in *Ranitomeya imitator*.** We extracted protein from *R. imitator* tadpole brains. Untreated protein (lane 1), protein incubated with protein phosphatase 1 (PP1) in dephosphorylation buffer overnight (lane 2), and protein incubated in dephosphorylation buffer without PP1 overnight (lane 3) were used. The untreated protein sample shows a single band ~37 kDA, the predicted weight. The sample treated with dephosphorylation buffer but not PP1 also had a faint band ~37 kDA. However, the sample incubated with PP1 overnight was not detected with the pS6 antibody, indicating that the used pS6 antibody is specific to only phosphorylated S6 ribosomal subunits in *R. imitator*.
