## Supplementary figures and images for "Tadpoles rely on mechanosensory stimuli for communication when visual capabilities are poor"

### Supplemental File 1

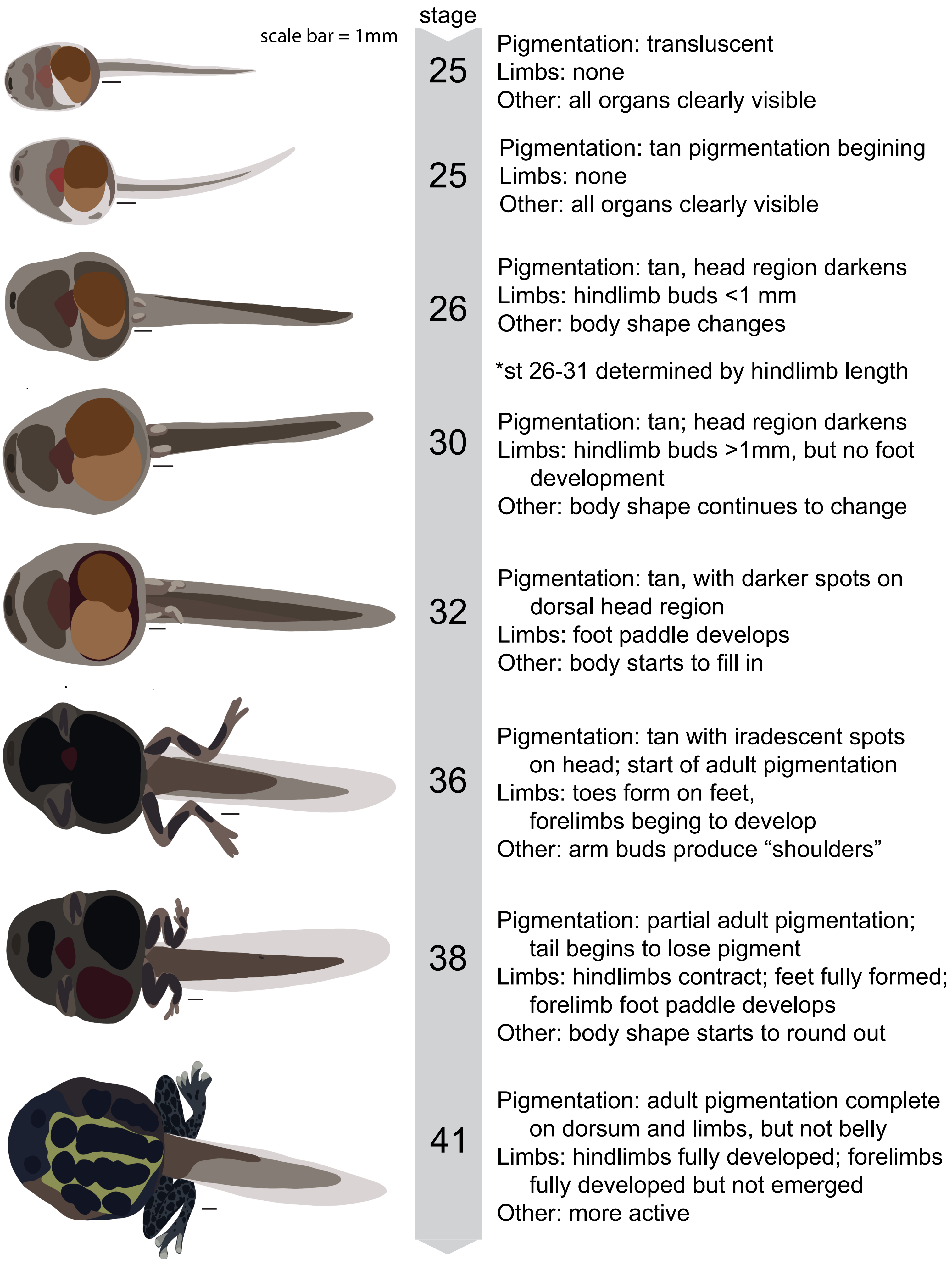
