## Supplemental File 3 for "Tadpoles rely on mechanosensory stimuli for communication when visual capabilities are poor"

#### R pipeline for analyzing phosphoTRAP data
#### Butler et al, 2022, Visual System Development in Social Poison Frog Tadpoles

### input file: count data corrected for read depth
### column 1 = gene IDs
### remaining columns = adjusted count data for input and ip samples


### load needed packages 
library(glmmTMB)
library(dplyr) 
library(stringr)
library(Biostrings)
library(ggplot2)

###### load and format data #####

### create list of column names (in correct order). Just here to double check importing correctly. 
conames <- list ("GeneID", "b1input", "b1ip", "b2input", "b2ip", "b3input", "b3ip", "nb1input", "nb1ip", 
                 "nb2input", "nb2ip", "nb3input", "nb3ip")

### load data
counts <- read.table(file="rimibeg_eye.gene.counts.matrix", header=TRUE, col.names = conames)

### remove the genes with all 0s
counts2 <- counts[!(counts$b1input=="0" & counts$b2input=="0" & counts$b3input=="0" & counts$b1ip=="0" & counts$b2ip=="0" & counts$b3ip=="0"),]
##leaves 25048 genes


########## running begging and non-begging separately ##########


########## for begging samples ##########

### separate begging and non-begging count data
### format it into a long table for analysis
counts_beg <- counts2[,c(1,3,2,5,4,7,6)]
B1 <- cbind(GeneID=counts_beg[,1], id="B1", IP=counts_beg[,2], IN=counts_beg[,3])
B2 <- cbind(GeneID=counts_beg[,1], id="B2", IP=counts_beg[,4], IN=counts_beg[,5])
B3 <- cbind(GeneID=counts_beg[,1], id="B3", IP=counts_beg[,6], IN=counts_beg[,7])

counts.beg.long<-as.data.frame(rbind(B1,B2,B3), stringsAsFactors = FALSE)
counts.beg.long.sort<- arrange(counts.beg.long, GeneID)  
counts.beg.long.sort$IP<-as.numeric(counts.beg.long.sort$IP)
counts.beg.long.sort$IN<-as.numeric(counts.beg.long.sort$IN)
colnames(counts.beg.long.sort)<- c("GeneID","Animal","ip.count","input.count")


###### compare IP to IN count data using paired t-tests (begging) #####
### data is in counts.beg.long.sort
### 3 samples per contig with 25048 contigs after removing zeros

### create an empty dataframe to populate
pairedt_beg_bnb_eye <- data.frame(GeneID=character(), t.value=numeric(), p.value=numeric(), stringsAsFactors = FALSE)

#run paired t-tests on all transcripts (begging only)
for (i in 1:25048)
{
  tempSamples <- counts.beg.long.sort[((3*i)-2):(3*i),]
  test<-t.test(tempSamples$ip.count, tempSamples$input.count, paired = TRUE)
  pairedt_beg_bnb_eye[i,] <- c(as.character(tempSamples[1,1]),test$statistic,test$p.value)
}


### check how many have a significant p-value
table(pairedt_beg_bnb_eye$p.value<0.05) #3187 transcripts where IP is different from INPUT, within begging only


###### calculate fold change for beg IP to IN (begging) #####

### let's do some reformatting fo calculating fold changes
row.names(counts2)<- counts2[,1] #move gene ID to row ID
counts2 <- counts2[,-1] #remove column 1 (gene ID)
View(counts2) #check to make sure works
counts3 <- counts2 + 1 #have to do this to create ratios

#separating out IP and IN within begging
counts_beg.ip <- counts3[,c(2,4,6)]
counts_beg.tot <- counts3[,c(1,3,5)]
ratios.beg <- counts_beg.ip/counts_beg.tot
logratiosbeg <- log2(ratios.beg)
logFC.beg <- data.frame("GeneID"=row.names(logratiosbeg),"logFC"=rowMeans(logratiosbeg))


###### add gene names #####
genenames <- read.table(file="RiTad_GeneNames.csv", header=TRUE, sep = ",") #load the gene names file
genenames$GeneID <- str_remove(genenames$GeneID, "_i1") #remove the isoform designation
genenames <- genenames[,-1] #removes column 1 which is blank for some reason...  


###### combine all into one file for plotting and saving #####
rimibeg_pairedT_eye_final <- merge(pairedt_beg_bnb_eye, logFC.beg, by="GeneID")
rimibeg_pairedT_eye_final <- merge(rimibeg_pairedT_eye_final, genenames, by="GeneID")

### for some reason it changes to characters, so force back to numeric
rimibeg_pairedT_eye_final$p.value <- as.numeric(rimibeg_pairedT_eye_final$p.value)

### first lets label each transcript as differentially expressed or not
rimibeg_pairedT_eye_final$diffexpressed <- "NO" #add NO to all transcripts, but we'll change the ones that are up and down in the next couple steps
rimibeg_pairedT_eye_final$diffexpressed[rimibeg_pairedT_eye_final$logFC > 0.6 & rimibeg_pairedT_eye_final$p.value < 0.05] <- "UP"
rimibeg_pairedT_eye_final$diffexpressed[rimibeg_pairedT_eye_final$logFC < -0.6 & rimibeg_pairedT_eye_final$p.value < 0.05] <- "DOWN"

### ggplot volcano plot
p <- ggplot(data=rimibeg_pairedT_eye_final, aes(x=logFC, y=-log10(p.value), col=diffexpressed)) + geom_point() + theme_minimal()
p2 <- p + geom_vline(xintercept=c(-0.6, 0.6), col="red") +
  geom_hline(yintercept=-log10(0.05), col="red")
mycolors <- c("blue", "red", "black")
names(mycolors) <- c("DOWN", "UP", "NO")
p3 <- p2 + scale_colour_manual(values = mycolors)
p3

### how DEGs are up or down, based on both fold change and p-value? 
table(rimibeg_pairedT_eye_final$diffexpressed=="UP") # = 83 enhanced in begging 
table(rimibeg_pairedT_eye_final$diffexpressed=="DOWN") # 2389 depleted in begging

### make a list of DEGs
rimibeg_beg_eye_DEGs_up <- rimibeg_pairedT_eye_final[(rimibeg_pairedT_eye_final$diffexpressed=="UP"),] 
rimibeg_beg_eye_DEGs_down <- rimibeg_pairedT_eye_final[(rimibeg_pairedT_eye_final$diffexpressed=="DOWN"),] 

### write out the files... 
write.csv(rimibeg_pairedT_eye_final, file="rimibeg_eye_beg_pairedT.csv")
write.csv(rimibeg_beg_eye_DEGs_up, file="rimibeg_eye_pairedT_beg_DEGs_up.csv")
write.csv(rimibeg_beg_eye_DEGs_down, file="rimibeg_eye_pairedT_beg_DEGs_down.csv")


########## now going to run it in the non-begging eye samples ##########


###### preparing the data #####

### create count file with only non-begging samples
counts_non <- counts2[,c(1,9,8,11,10,13,12)]

### remove transcripts where all counts are 0
counts_non <- counts_non[!(counts_non$nb1ip=="0" & counts_non$nb1input=="0" & counts_non$nb2ip=="0" & counts_non$nb2input=="0" & counts_non$nb3ip=="0" & counts_non$nb3input=="0"),]

### format a long data file for analysis
NB1 <- cbind(GeneID=counts_non[,1], id="NB1", IP=counts_non[,2], IN=counts_non[,3])
NB2 <- cbind(GeneID=counts_non[,1], id="NB2", IP=counts_non[,4], IN=counts_non[,5])
NB3 <- cbind(GeneID=counts_non[,1], id="NB3", IP=counts_non[,6], IN=counts_non[,7])

counts.non.long<-as.data.frame(rbind(NB1,NB2,NB3), stringsAsFactors = FALSE)
counts.non.long.sort<- arrange(counts.non.long, GeneID)  
counts.non.long.sort$IP<-as.numeric(counts.non.long.sort$IP)
counts.non.long.sort$IN<-as.numeric(counts.non.long.sort$IN)
colnames(counts.non.long.sort)<- c("GeneID","Animal","ip.count","input.count")


###### compate IP and IN counts using paried t-tests (non-begging) #####
### data is in counts.non.long.sort
### 3 samples per contig with 25084 contigs after removing zeros 

### create an empty dataframe
pairedt_non_bnb_eye <- data.frame(GeneID=character(), t.value=numeric(), p.value=numeric(), stringsAsFactors = FALSE)

### run the paired t-tests
for (i in 1:25048)
{  
  skip_to_next <- FALSE
  
  tryCatch(
    {tempSamples <- counts.non.long.sort[((3*i)-2):(3*i),]
    test<-t.test(tempSamples$ip.count, tempSamples$input.count, paired = TRUE)
    pairedt_non_bnb_eye[i,] <- c(as.character(tempSamples[1,1]),test$statistic,test$p.value)}, 
    error = function(e) {skip_to_next <<- TRUE})
  if(skip_to_next) { next }
  
}

#lets check how many transcripts are sig. different
table(pairedt_non_bnb_eye$p.value<0.05) #631 significant p-values


###### calculate fold change for beg IP to IN (non-begging) ####

### some reformatting for this
row.names(counts2)<- counts2[,1] #move gene ID to row ID
counts2 <- counts2[,-1] #remove column 1 (gene ID)
View(counts2) #check to make sure works
counts3 <- counts2 + 1 #have to do this for ratios 

counts_non.ip <- counts3[,c(8,10,12)]
counts_non.tot <- counts3[,c(7,9,11)]
ratios.non <- counts_non.ip/counts_non.tot
logratiosnon <- log2(ratios.non)
logFC.non <- data.frame("GeneID"=row.names(logratiosnon),"logFC"=rowMeans(logratiosnon))


###### add gene names #####
genenames <- read.table(file="RiTad_GeneNames.csv", header=TRUE, sep = ",") #load the gene names file
genenames$GeneID <- str_remove(genenames$GeneID, "_i1") #remove the isoform designation
genenames <- genenames[,-1] #removes column 1 which is blank for some reason  


###### combine all into one file for plotting and saving it #####
rimibeg_pairedT_eye_non_final <- merge(pairedt_non_bnb_eye, logFC.non, by="GeneID")
rimibeg_pairedT_eye_non_final <- merge(rimibeg_pairedT_eye_non_final, genenames, by="GeneID")

### for some reason it changes to characters, so force back to numeric
rimibeg_pairedT_eye_non_final$p.value <- as.numeric(rimibeg_pairedT_eye_non_final$p.value)

### first lets label each transcript as differentially expressed or n
rimibeg_pairedT_eye_non_final$diffexpressed <- "NO"
rimibeg_pairedT_eye_non_final$diffexpressed[rimibeg_pairedT_eye_non_final$logFC > 0.6 & rimibeg_pairedT_eye_non_final$p.value < 0.05] <- "UP"
rimibeg_pairedT_eye_non_final$diffexpressed[rimibeg_pairedT_eye_non_final$logFC < -0.6 & rimibeg_pairedT_eye_non_final$p.value < 0.05] <- "DOWN"

### ggplot vlcano plot
q <- ggplot(data=rimibeg_pairedT_eye_non_final, aes(x=logFC, y=-log10(p.value), col=diffexpressed)) + geom_point() + theme_minimal()
q2 <- q + geom_vline(xintercept=c(-0.6, 0.6), col="red") +
  geom_hline(yintercept=-log10(0.05), col="red")
mycolors <- c("blue", "red", "black")
names(mycolors) <- c("DOWN", "UP", "NO")
q3 <- q2 + scale_colour_manual(values = mycolors)
q3

### how DEGs are up or down, based on both fold change and p-value? 
table(rimibeg_pairedT_eye_non_final$diffexpressed=="UP") # = 60 enhanced in non-begging 
table(rimibeg_pairedT_eye_non_final$diffexpressed=="DOWN") # = 443 depleted in non-begging

### make a list of DEGs
rimibeg_eye_non_DEGs_up <- rimibeg_pairedT_eye_non_final[(rimibeg_pairedT_eye_non_final$diffexpressed=="UP"),] 
rimibeg_eye_non_DEGs_down <- rimibeg_pairedT_eye_non_final[(rimibeg_pairedT_eye_non_final$diffexpressed=="DOWN"),] 

### write out the files... 
write.csv(rimibeg_pairedT_eye_non_final, file="rimibeg_brain_pairedT.csv")
write.csv(rimibeg_eye_non_DEGs_up, file="rimibeg_brain_pairedT_non_DEGs_up.csv")
write.csv(rimibeg_eye_non_DEGs_down, file="rimibeg_brain_pairedT_non_DEGs_down.csv")
