## Supplemental File 7 for "Tadpoles rely on mechanosensory stimuli for communication when visual capabilities are poor"

### *Ranitomeya imitator* Tadpole Ethogram

| Behavior | | Classification | Description |
|---|---|---|---|
| Swimming | 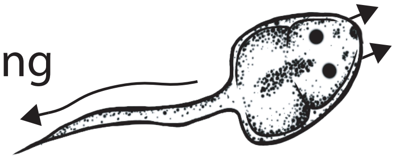 | general | Involves several tail movements to create a continuous movement covering distance around the behavior arena. |
| Movement | 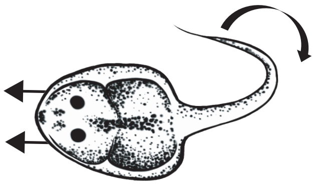 | general | Single tail flick propels tadpole to a new location in arena, but followed by at least 1 second of no additional movement. |
| Roll (side view) | 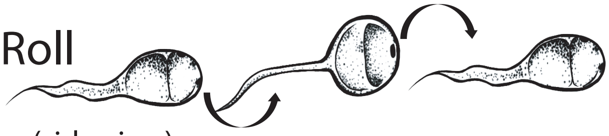 | general | Tadpole rolls ~90$^{o}$ horizontally, exposing their ventral surface to the side. |
| Wiggling | 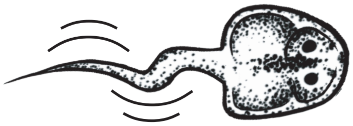 | general/ affiliative | Tail curls back and forth to create a wiggling motion that is not vibrational. Often, but not always, directed at an adult. |
| Begging | 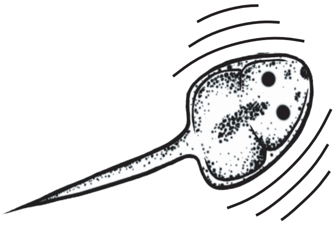 | affiliative | Rapid whole body vibration that creates an obvious water distrubance. Tail is straight, but often at an angle to the body. |
| Nip | 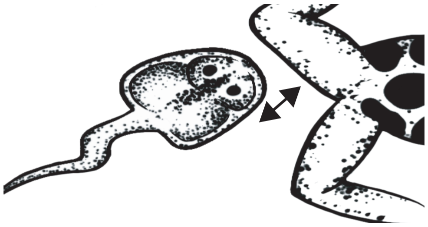 | affiliative | Tadpole mouths at adult, tpically directed towards the legs or back end of the adult. |
| Bite/Ram | 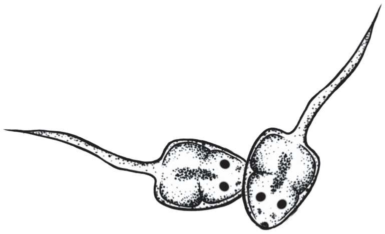 | aggressive | One tadpole, with mouth open (bite) or closed (ram), hits/bites the body of another tadpole. Occassionally, tadpole will maintain bite and thrash around (bite w/ thrashing). |
